# An R Implementation of Tumor-Stroma-Immune Transcriptome Deconvolution Pipeline using DeMixT

**DOI:** 10.1101/566075

**Authors:** Shaolong Cao, Zeya Wang, Fan Gao, Jingxiao Chen, Feng Zhang, Daniel E. Frigo, Eleni Efstathiou, Scott Kopetz, Wenyi Wang

**Affiliations:** Department of Bioinformatics and Computational Biology, The University of Texas MD Anderson Cancer Center, Houston, TX, USA; Department of Statistics, Rice University, Houston, TX, USA; UT Health School of Public Health, Houston, TX, USA; Department of Cancer Systems Imaging, The University of Texas MD Anderson Cancer Center, Houston, TX, USA; Department of Genitourinary Medical Oncology, The University of Texas MD Anderson Cancer Center, Houston, TX, USA; Department of Gastrointestinal (GI) Medical Oncology, The University of Texas MD Anderson Cancer Center, Houston, TX, USA

**Keywords:** Tumor heterogeneity, Tumor-stroma-immune interaction, TCGA RNA-seq deconvolution, Parallel computing, Adaptive numerical integration

## Abstract

The deconvolution of transcriptomic data from heterogeneous tissues in cancer studies remains challenging. Available software faces difficulties for accurately estimating both component-specific proportions and expression profiles for individual samples. To address these challenges, we present a new R-implementation pipeline for the more accurate and efficient transcriptome deconvolution of high dimensional data from mixtures of more than two components. The pipeline utilizes the computationally efficient DeMixT R-package with OpenMP and additional cancer-specific biological information to perform three-component deconvolution without requiring data from the immune profiles. It enables a wide application of DeMixT to gene expression datasets available from cancer consortium such as the Cancer Genome Atlas (TCGA) projects, where, other than the mixed tumor samples, a handful of normal samples are profiled in multiple cancer types. We have applied this pipeline to two TCGA datasets in colorectal adenocarcinoma (COAD) and prostate adenocarcinoma (PRAD). In COAD, we found varying distributions of immune proportions across the Consensus Molecular Subtypes, from the highest to the lowest being CMS1, CMS3, CMS4 and CMS2. In PRAD, we found the immune proportions are associated with progression-free survival (p<0.01) and negatively correlated with Gleason scores (p<0.001). Our DeMixT-centered analysis protocol opens up new opportunities to investigate the tumor-stroma-immune microenvironment, by providing both proportions and component-specific expressions, and thus better define the underlying biology of cancer progression.

Availability and implementation: An R package, scripts and data are available: https://github.com/wwylab/DeMixTallmaterials.

## Background

Solid tumor tissue samples often consist of more than one distinct tissue compartment. For example, the tumor-immune cell contexture has been anatomically demonstrated to primarily consist of a tumor core, lymphocytes and other non-cancerous cells ^1,2^. Thus, cellular heterogeneity adds confounding complexity to genomic analyses in cancer studies, complicating the evaluation of individualized clinical outcomes. Recently, several computational approaches have been proposed to help understand the heterogeneity of tumor tissues. These approaches are rationalized to circumvent the high expense and long time from previous experimental approaches such as laser-capture microdissection (LCM) and cell sorting for deconvolution. Most commonly available computational methods using high-throughput expression data faces distinct challenges. Some are limited to two distinct components, without consideration of immune cells^3,4^. Others require knowledge of cell-type-specific gene lists, i.e., reference genes^5-8^. In addition, some do not account for sample variances within each component, which can result in large biases in the estimated mixing proportions and mean expressions^9^. We recently proposed a statistical modeling framework, DeMixT, for jointly estimating the proportions and component-specific and sample-specific gene expression levels with more than two components^10^.

To perform deconvolution, the DeMixT method requires expression data from *p*-1 components, where *p* is either 2 or 3, the total number of components for deconvolution. The Cancer Genome Atlas (TCGA) projects have generated high-throughput RNAseq data in over 11,000 tumor samples across 33 cancer types, among which 16 have data from normal tissue samples. None of the cancer types in TCGA has data from expression profiles of the actual immune component. Previously, we successfully ran DeMixT 3-component deconvolution on TCGA head-neck squamous cell carcinoma (HNSC) dataset, using biological information such as the human papillomavirus (HPV) status^10^. To overcome the likely persisting issue of lacking immune profiles, we have developed a general analysis pipeline to perform three component deconvolution, i.e., stroma, immune and tumor, using datasets from TCGA or studies of a similar design. Other persisting issues in deconvolution include computational efficiency and accuracy, due to the high dimensional parameter space we are interrogating. Here, we describe top-down, the implementation and results of the new pipeline, from the pipeline setup, to the DeMixT R package with parallel computing, then to a new adaptive numerical integration approach for the core repetitive calculation inside DeMixT.

## Implementation

### A three-component deconvolution pipeline that requires only reference profiles from the stromal component

This pipeline is applicable to the TCGA datasets where the gene expression data from mixed tumor samples and normal samples are available but no immune profiles are provided. As shown in **Figure 1,** we first select a set of tumor samples that are known to be high in tumor content (e.g., DeMixT 2-component proportion > 85%) to obtain the tumor-specific expression profile. We then select a set of tumor samples that are known to be high in immune content (clinically annotated) to use the tumor-specific expression profile and the normal profile to obtain the immune-specific expression profile. This immune profile is then used together with the normal profile and the full set of mixed tumor samples as input data of DeMixT to obtain the final results.

**Figure 1:**
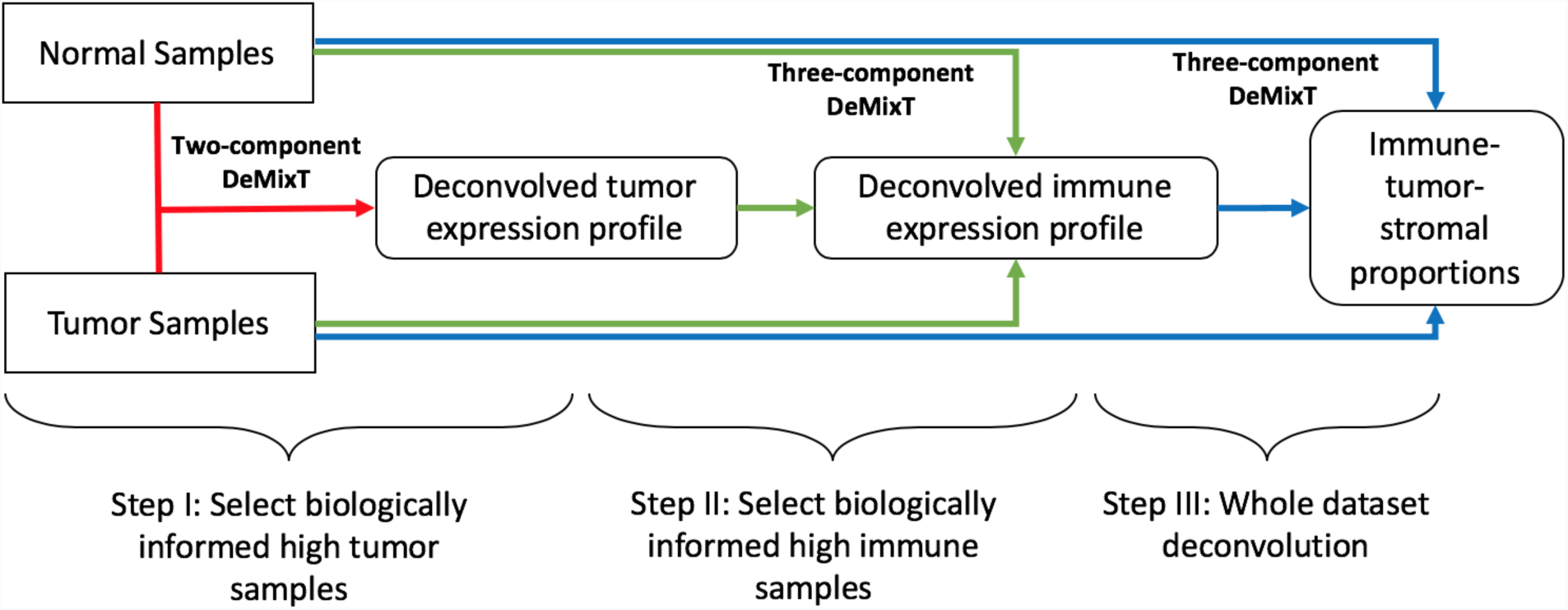
Three-component deconvolution pipeline using DeMixT. Normal samples are the reference samples and tumor samples are the observed mixed tumor samples.

We have developed the corresponding pipelines for colon adenocarcinoma (COAD) and prostate adenocarcinoma (PRAD) in TCGA data with immune indicators, microsatellite instability levels (MSI)^11^, and mutation burden, respectively. The mutation burden for each PRAD sample is calculated based on the TCGA MC3 consensus mutation callers^12^.

### DeMixT

Our model expresses the observed gene expressions from each gene and each mixed tumor sample as a weighted linear combination of unobserved raw expressions from the components, which each follow a log_2_-normal distribution and is weighted by its corresponding proportion. The current maximum number of components is three. One component can be assumed unknown, while the distributions of the remaining components need to be estimated from reference samples. DeMixT adopts an approach of iterated conditional modes (ICM) to maximize the full likelihood^13^. We further used a gene-set-based component merging approach (GSCM) to reduce the bias of proportion estimation for three-component deconvolution^10^.

### DeMixT R package

A main function DeMixT is designed to finish the whole pipeline of deconvolution for two or three components. The DeMixT.S1 function is designed for the first step to estimate the proportions of all mixed samples for each component. The DeMixT.S2 function implements the second step to estimate the component-specific deconvolved expressions of individual mixed samples for a given set of genes. For more flexible manipulation of features in DeMixT, a function Optimum.KernelC is provided to modify values of function running parameters used in the deconvolution steps. R packages DeMixT_0.2 and DeMixT_0.2.1 are the latest versions. DeMixT_0.2 is designed for a computer that has OpenMP (strongly recommended for computational efficiency). DeMixT_0.2.1 is designed for computers that do not have OpenMP. OpenMP enables parallel computing on a single computer by taking advantage of the multiple cores available on modern CPUs. The ICM framework further enables parallel computing, which helps compensate for the extensive computing time used in the repeated numerical double integrations.

### Adaptive numerical integration underlying the DeMixT likelihood calculation

The fundamental computing block of DeMixT is to estimate the likelihood of each gene and sample. Assume the hidden expression profile of each component are *N*_*1*_, *N*_*2*_ and *T*, the observed expression is *Y*, the linear additivity for component-specific expressions under raw measured data follows *Y*_*ig;*_ = π_*1,i*_*N*_*1,ig*_ + π_*2,i*_*N*_*2ig*_ + (*1* – π_*1,i*_ – π_*2,i*_)*T*_*ig*_, where *g* = *1,2*, …, *G* and *i* = *1,2*, …, *S* stand for gene and sample respectively. For hidden expression level of each component, we assume they follow the log2-normal distribution. 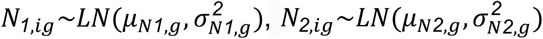 and 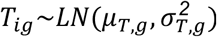. For a given gene and sample, the likelihood can be calculated based on the following^10^:

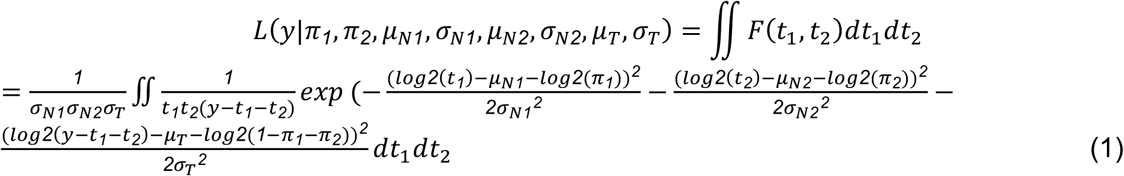

Where *F*(*t*_1_, *t*_2_) is probability density function, *t*_1_ = π_*1, i*_*N*_*1,ig*_ and *t*_2_ = π_*2,i*_*N*_*2,ig*_ are two random variables that follow log-normal distributions. Maximum likelihood estimators are then derived for the unknown parameters μ_*T*_, σ_*T*_, π_*1*_, π_*2*_ in equation (1). DeMixT uses ICM algorithm to update proportion parameters π_*1,i*_, π_*2,i*_ and gene parameters 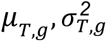 iteratively. We reduced the optimization problem to a one-dimensional search in two-dimensional space for each pair of parameters, using golden section search with parabolic interpolation. Assuming for the estimation of each parameter, it takes K steps to reach the maximum likelihood in each iteration, and N full iterations of to reach the stop criterion, we can roughly estimate that the numerical integration needs to be repeated O(K*N*S*G) times. For example, the integration needs to be repeated about 600 million times for a gene expression dataset with 200 tumor samples and 15,000 genes, when N=20 and K=10. To ensure computational efficiency and accuracy, we have devised a new adaptive numerical integration approach.

The triangle in **Figure 2A** represents domain of (*t*_1_, *t*_2_) for the probability function *F*(*t*_1_, *t*_2_), which is shown as a 3-D peak in **Figures 2B**, **2C** and **2D**. When *F*(*t*_1_, *t*_2_) falls in the boundary region of the domain space of (*t*_1_, *t*_2_), as shown in **Figure 2A** in yellow, blue and grey, it is peaked (>75% of total peak volume) in a small region. When *F*(*t*_1_, *t*_2_) are centered, as shown in **Figure 2A** in red, it is more spread out in that region, justifying for an even grid summation. The regular two-dimensional integration, i.e., rectangle method, starts with dichotomizing the integration domain (triangle) into even size grid (square), then summing up all integration values across all grids. Instead of evenly separating the grids within the integration domain, our new adaptive integration re-distributes more grids (maintaining the same total amount of grids for stable computing time, default at 3,600) near the mode of probability density when it sits inside the boundary area: yellow, blue and grey (**Figure 2A**). By doing this, the adaptive integration is able to achieve better accuracy using the same number of grids. Further details of the adaptive integration algorithm are provided in the **Supplementary Materials**.

**Figure 2:**
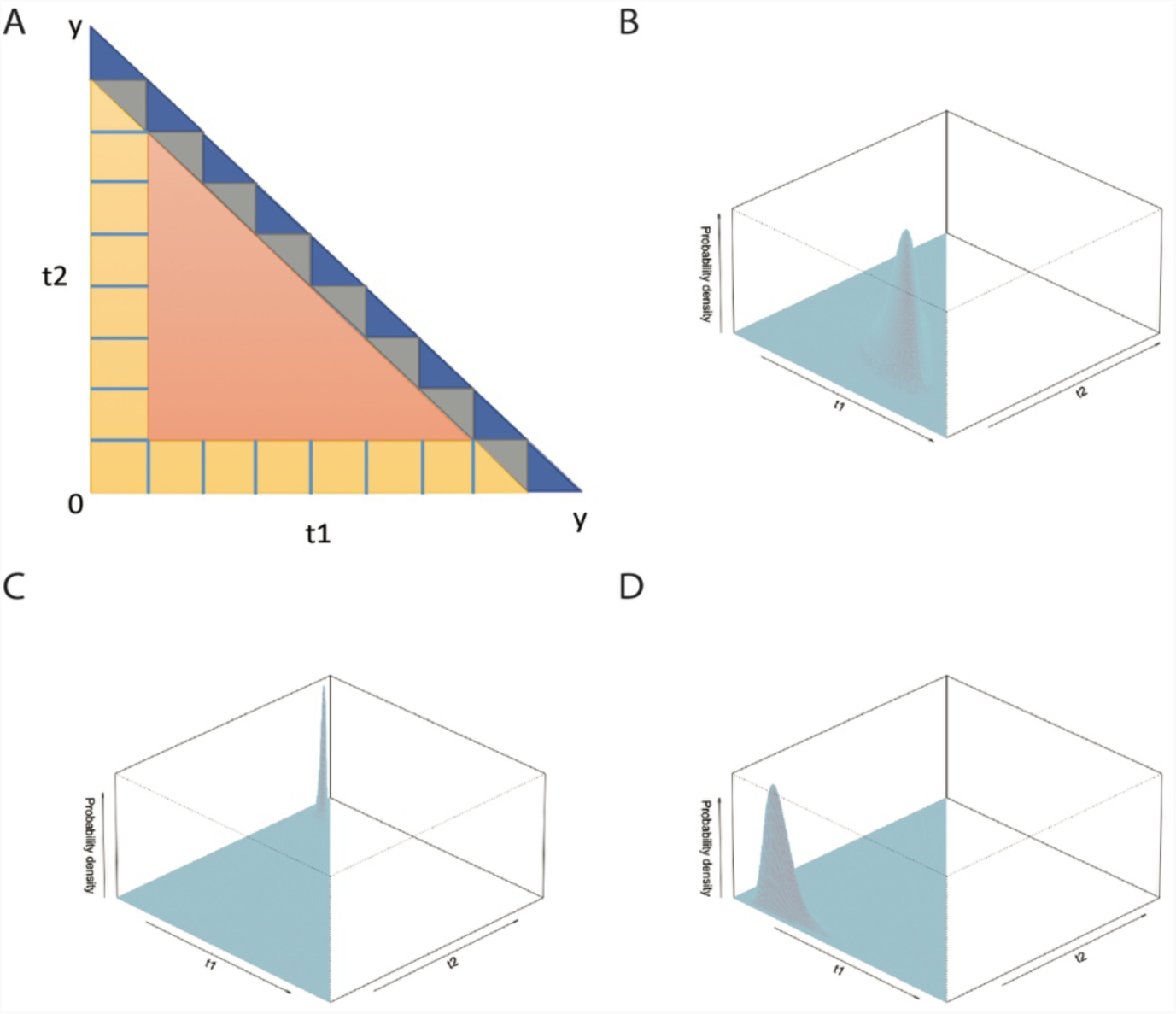
Adaptive numerical integration of the DeMixT likelihood function. **A.** The integration triangle region of the full likelihood function for three components are dichotomized into four regions (represented as four colors). We first identify the location of the peak of probability density and then adaptively increase the number of bins near the peak area. An adaptive integration strategy is determined by which region (color) the peak falls into. **B,C,D.** Three examples of the three-dimensional surface plot of the integral function for each sample and gene. Z axis represents the probability density. The surface in general has a unique peak. The peak in **B** sits in the red area. The peak in **C** sits in the blue area. The peak in **D** sits in the yellow area.

## Results

Different from previous computational deconvolution methods, DeMixT provides a comprehensive view of tumor-stroma-immune transcriptional dynamics, as compared to methods that address only immune subtypes within the immune component, in each tumor sample. Although advanced in theoretical methods, it is still challenging to differentiate tumor and immune levels among cancer patients simply based on statistical models without cancer-specific biological guidance. We have now designed a general three-component deconvolution pipeline to allow for the application of DeMixT to TCGA-type gene expression datasets where only data from the mixed tumors and normal samples are available. We further incorporated cancer-type-specific biological information into the pipeline for the transcriptome deconvolution in PRAD and COAD. The deconvolved proportions and component-specific expression matrices (for each sample and each gene) for these two cancer types are available for download at http://bioinformatics.mdanderson.edu/public-software/demixt.

### TCGA Prostate Adenocarcinoma (PRAD) dataset^14^

We generated tumor-stroma-immune proportions for PRAD by selecting tumor samples with tumor purity higher than 85% in a two-component deconvolution to create the tumor profile, and then selecting high immune samples guided by mutation burden (total number of mutations) larger than 100 to build the immune profile^15^. Here the mutation burden is calculated based on the consensus mutation calls^12^ made on the whole-exome sequencing data generated from the same sample. Across 295 samples, the immune proportions have a median of 29% and a median absolute deviation (MAD) of 16%; the tumor proportions have a median of 56% and a MAD of 24%; and stroma proportions have a median of 12% and a MAD of 9% (**Figure 3A**). With these proportions available, we compared the immune proportions with clinical annotations of the patients samples. Interestingly, we saw significantly different immune proportions across patients with different Gleason scores (6, 7 and >7, p<0.001, **Figure 3C**), with high Gleason scores presenting the lowest immune proportions. Gleason score is a grading system developed by pathologist that is widely used to evaluate the aggressiveness of prostate cancer clinically^16^. Patients graded with higher Gleason scores (>7) have more aggressive tumors. We further divided the patients into two equal-sized groups: high immune (>29%) and low immune (<29%), and found statistically significant differences in progression-free survival (PRS) outcome between the two groups, after adjusting for age and sex (p=0.002, **Figure 3D**). PFS is the period between the date of the first cancer diagnosis and the date of a new occurrence of a tumor event. It has been shown to be a good surrogate for cancer specific survival in prostate cancer^17^, given the long latency for prostate cancer and relative limited follow-up time of this more recent clinical cohort. Importantly, patients with high estimated immune proportions had significantly higher progression-free probability than patients with low estimated immune proportions.

**Figure 3:**
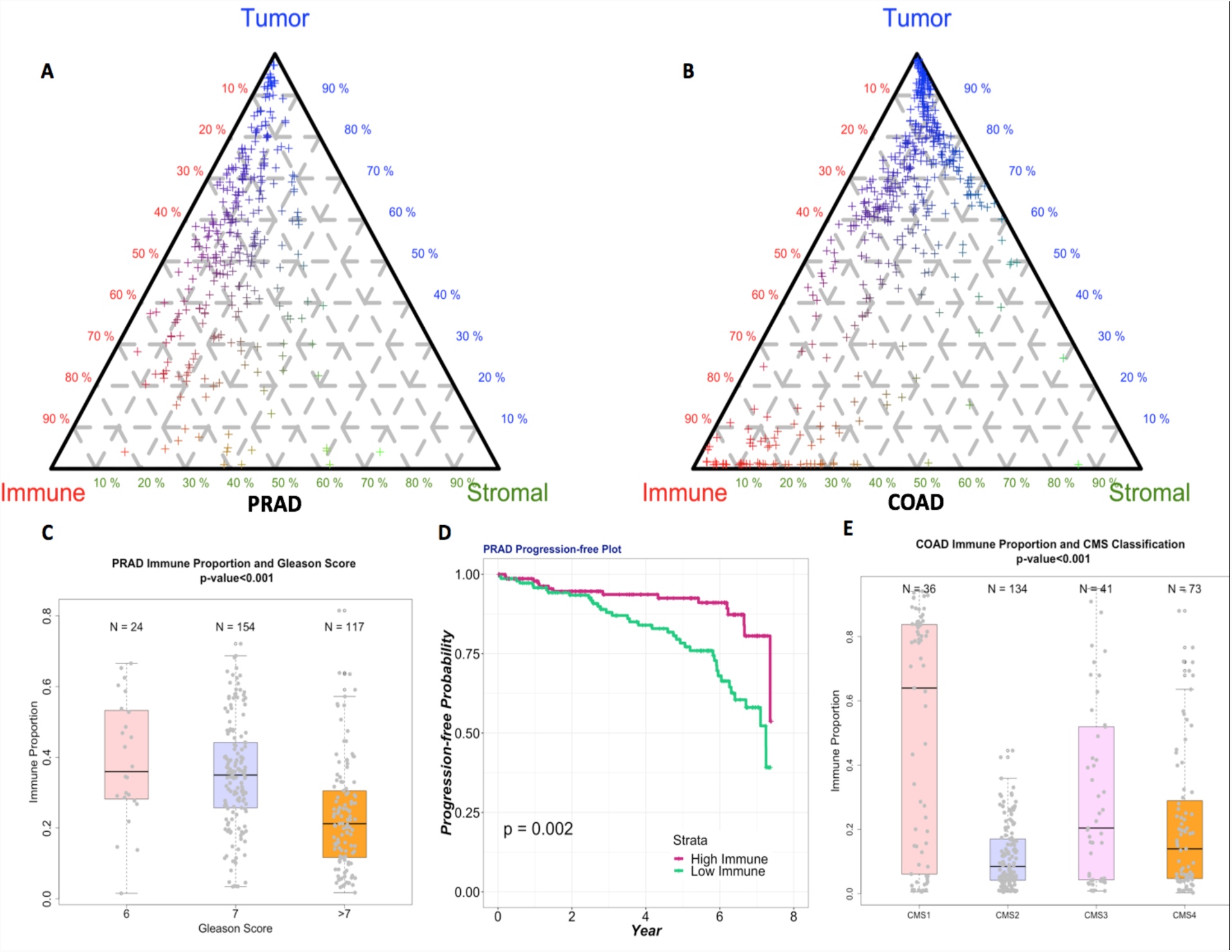
Three-component deconvolution results and validation. **A&B**. Tri-plots of estimated proportions (%) of the tumor component (top), the immune component (bottom left), and the stromal component (bottom right) in the PRAD and COAD data. Points closer to a component’s vertex suggest higher proportion for the corresponding component, whose quantity equals the distance between the side opposite the vertex and a parallel line (illustrated as dashed gray lines for the multiples of 10th percentile) that a point is sitting on. **C**. Boxplots of estimated immune proportions (%) in PRAD patients with the corresponding Gleason scores. **D**. Kaplan-Meier estimator in the PRAD based on progression-free survivals with p-value from a coxPH model. **E.** Boxplots of estimated immune proportions (%) with corresponding CMS classification. Sample sizes of each category are on top of each boxplot and the p-value is calculated with Krustal-Wallis test.

### TCGA Colorectal Adenocarcinoma (COAD) dataset^18^

Similarly, the tumor-stroma-immune proportions for COAD were generated by selecting high tumor samples with tumor purity higher than 90% (this cutoff is also cancer-type specific), and then selecting samples who were of high microsatellite instability^11^ as high immune samples. Out of the 442 patient samples, we observed the median of tumor proportion is 69% with MAD 22%, the median of immune proportion is 15% with MAD 17% and the median of stroma proportion is 11% with MAD 8% (**Figure 3B**). In colorectal cancer, recent studies on the consensus molecular subtypes (CMS) classification system have identified four consensus molecular subtypes, termed CMS1-4^19^. The four distinct CMS subtypes, classified based on 2000 gene expression profiles in the primary tumor, can have different survival outcomes. CMS1 samples are hypermutated with the least prevalence of somatic-number alterations (SCNs) and are expected to have the highest immune infiltration and activation compared to the other CMSs. CMS2 is the opposite case of CMS1 with the least mutation counts but the most SCNs and are expected to be least immune infiltrated. Without having such prior knowledge, our immune proportions demonstrated a similar distribution across CMS subtypes (p-value<0.001, **Figure 3E**), supporting the utility of our pipeline to accurately estimate tumor-stroma-immune proportions.

To evaluate the performance of the proposed adaptive numerical integration strategy, we simulated 50 mixed samples with 1000 genes. The proportion of π_1_ T component was generated evenly between 0.15 and 0.85 and the mixing proportions of *N*_1_ and *N*_2_ components are randomly generated from a uniform distribution such that π_1_ + π_2_ + π_*T*_ = 1 For each gene, the expression of *N*_1_, *N*_2_ and *T* were sampled independently by 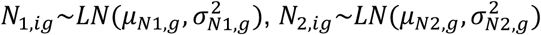 and 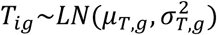 where μ_*N1,g*_, μ_*N*2*,g*_, μ_*T,g*_∼*N*(7,3.5), σ_*N1,g*_ = σ_*N2,g*_ = σ_*T,g*_ = 0.5. Finally, the observed was calculated by *Y*_*ig*_ = π_1,*i*_*N*_1,*ig*_ + π_2*i*_*N*_2,*ig*_ + (1 – π _1*i*_ – π_2*i*_)*T*_*ig*_. We compared the proposed adaptive integration with the regular numerical integration. The regular numerical integration was implemented by evenly distributed bins on the triangle integration region (**Figure 2A**). To ensure a fair comparison, we limited the number of bins to ensure the same numbers between regular and adaptive integration, e.g., the total bins were fixed to 3,600 for each sample and gene. We used concordance correlation coefficient (CCC) and Root Mean Squared Error (RMSE) to compare between the estimated μ_*T,g*_ π_*T*_ and true values using the two approaches for the likelihood calculation, respectively. Higher values in CCC (maximum is 1) and lower values in RMSE (minimum is 0) suggest better performances. As shown in **Table 1**, both criteria demonstrated a highly improved performance for parameter estimation when we use the adaptive numerical integration.

**Table 1.** Adaptive and regular integration comparison

|  | RMSE | CCC |
| --- | --- | --- |
| $\mu_{T,g}$ (adaptive) | 0.833 | 0.896 |
| $\mu_{T,g}$ (regular) | 1.663 | 0.718 |
| $\pi_T$ (adaptive) | 0.053 | 0.951 |
| $\pi_T$ (regular) | 0.097 | 0.851 |

Our R package is maintained on GitHub (wwylab/DeMixTallmaterials). Parallel computing through integrating utilities like OpenMP is essential for improving the efficiency of running DeMixT on large-scale cancer datasets, such as those from TCGA. On a PC with a 3.07-GHz Intel Xeon processor with 20 cores, DeMixT takes 14 min to complete the full three-component deconvolution task for 50 samples and 500 genes, while it takes about 16 hours without OpenMP.

We also tested on a TCGA BLCA dataset with 408 samples and 6,804 genes. The three-component deconvolution of DeMixT takes about 35 hours on a 5-core PC with 3.07-GHz Intel Xeon CPU, as compared to just 3.5 hours on a 56-core server with 2.60-GHz Intel Xeon Gold 6132 CPU. The DeMixT source files are compatible with Windows, Linux and MacOS. The OpenMP is readily installed in most Linux servers and Windows PCs. We have received user feedbacks that MacOS systems can have difficulties in the installation of OpenMP-enabled DeMixT R package. To help users resolve the potential installation issues we have provided a guideline about how to install OpenMP (http://bioinformatics.mdanderson.org/public-software/demixt) and a link to the OpenMP official website (http://www.openmp.org/).

## Discussion

We present an R implementation of three-component transcriptome deconvolution pipeline that is built on our R package DeMixT_0.2 with more practical requirement for the input data, i.e., only mixed tumor samples and normal samples. These types of datasets are available from large consortium studies such as TCGA. The output of our pipeline is unique as we provide a set of tumor-stroma-immune proportions that sum up to 1, providing a first standardized view of the three-component microenvironment, as well as component-specific expression levels. To improve the computational efficiency and accuracy of DeMixT, we have incorporated an adaptive numerical integration approach to solve the computation bottleneck of DeMixT, the likelihood calculation. DeMixT enables parallel computing through OpenMP, which is readily available on most Linux server systems. The runtime of DeMixT is O(1/number of cores). We therefore strongly recommend running the entire pipeline including DeMixT in the parallel computing mode on servers as compared to personal computers, in order to gain substantial reduction in runtime.

Cancer-specific, three-component deconvolution pipelines for other cancer types can be adapted from our general pipeline in a similar fashion. We consulted with and worked closely with physician scientists for reliable and benchmarked biological information in order to perform meaningful deconvolution in prostate and colorectal cancer. We therefore present this pipeline in a general term, expecting substantial biological input and effort to communicate across disciplines for further cancer-specific application, rather than an assumption-based-methods-guided analysis, e.g. using another deconvolution method as pseudo-truth. In our pipeline, we assumed the tumor-profiles derived from samples with high tumor content are representative of those in samples with high immune content. We also assumed that the immune profiles from samples with high immune content are representative across all samples. These assumptions will be further evaluated after we collect more biological information for deconvolution in future study, as it is a mostly unexplored territory. Future versions of the software will consider an extension with a hierarchical model for immune subpopulations that will allow deconvolution for more than three components and include dynamic immune components.

