## Supplementary Materials. for "An R Implementation of Tumor-Stroma-Immune Transcriptome Deconvolution Pipeline using DeMixT"

***An adaptive integration algorithm***

Our goal is to calculate the following likelihood of gene *g* and sample *i*.

$$L(y|\pi_1, \pi_2, \mu_{N1}, \sigma_{N1}, \mu_{N2}, \sigma_{N2}, \mu_T, \sigma_T) = \iint F(t_1, t_2) dt_1 dt_2$$

$$= \frac{1}{\sigma_{N1}\sigma_{N2}\sigma_T} \iint \frac{1}{t_1 t_2 (y - t_1 - t_2)} exp\,(-\frac{(log2(t_1) - \mu_{N1} - log2(\pi_1))^2}{2\sigma_{N1}{}^2}$$
$$- \frac{(log2(t_2) - \mu_{N2} - log2(\pi_2))^2}{2\sigma_{N2}^2}$$
$$- \frac{(log2(y - t_1 - t_2) - \mu_T - log2(1 - \pi_1 - \pi_2))^2}{2\sigma_T{}^2} dt_1 dt_2$$

Notation:

y $= Y_{ig}$: observed mixed expression for sample *i* and gene *g*, where $Y_{ig} = \pi_1 N_{1,ig} + \pi_2 N_{2,ig} + (1 - \pi_1 - \pi_2) T_{ig}$

$t_1 = \pi_1 N_{1,ig}$: where $N_{1,ig}$ is the hidden expression of $N_1$-component for sample *i* and gene *g*

$t_2 = \pi_2 N_{2,ig}$: where $N_{2,ig}$ is the hidden expression of $N_2$-component for sample *i* and gene *g*

$B_1$: number of bins for $t_1$ and $t_2$, e.g., 60 for regular integration or 30 for adaptive integration, depends on the location of the mode of the integral function $F(t_1, t_2)$

$B_2$: number of bins for small square *Sp* or small triangle *Tp*, e.g., 50

<u>*Rectangle integration method*</u>. For a given sample *i* and gene *g*, we first start from a basic grid separation. The idea is to cover the triangle, i.e., domain of integral function $F(t_1, t_2)$, using evenly sized small squares and small triangles along the diagonal. For example, we put a total of $B_1$ bins, evenly distributed, from 0 to *y* for $t_1$ axis, by drawing vertical lines to the $t_1$ axis. We then repeat the same strategy for the $t_2$ axis by drawing horizontal lines. The intersection of the vertical and horizontal lines will form the small squares as referred to in the main text. The triangle integration

region is dichotomized into $\frac{B_1(B_1-1)}{2}$ small squares and $B_1$ small triangles. The integration of equation (1) can be easily calculated by summing up all values of small squares: $\frac{F(x_1,x_2)*y^2}{B_1^2}$ and small triangles: $\frac{F(x_1,x_2)*y^2}{2B_1^2}$, where $(x_1, x_2)$ stands for the center position of each small square or small triangle.

*Step 1*. We first find the location of the peak of the integral function, i.e., $argmax\{F(t_1, t_2)\}$ using golden section search with parabolic interpolation. We determine which small square (as defined by the vertical and horizontal lines described above) contains the peak.

*Step 2A*. If the peak falls in the red area in **Figure 2A**, we set $B_1 = 60$ and simply use the rectangle integration method described above, as it is not necessary to perform adaptive integration.

*Step 2B*. If the peak falls in one of the small squares, denoted as *Sp*, in the yellow area in **Figure 2A**. In this case, the peak of the integral function is sharper and the contribution of the *Sp* square to the integration can be much larger as compared to other small squares. So we will allocate more computational resources to *Sp*, and merge other small squares to reduce their corresponding computation. We evenly divide up the small square *Sp* into $B_2 * B_2$ tiny squares. Then the integration value within the *Sp* can be calculated using rectangle integration method described above. Finally, adding up the integration of *Sp* with sums of other merged squares gives the integration result. In this case, we may set $B_1 = 30$ and $B_2 = 50$, for example, to maintain similar overall computating cost as step 2A.

*Step 2C.* If the peak falls in one of the small triangles, denoted as *Tp*, in the blue area in **Figure 2A**. We then separate the small triangle *Tp* into $\frac{B_2*(B_2-1)}{2}$ tiny squares plus $B_2$ tiny triangles and sum up the value of each tiny square using rectangle integration. Following the same procedure described in step 2B, the final results will be the integration inside *Tp* plus all other squares. In this case, we also set $B_1 = 30$ and $B_2 = 50$ for the same rationale.
